# Real-Time GPU-Accelerated Digital Heart Twin: Integrating Bidirectional Interactions Between Living Optogenetic Monolayers and Computational Simulations

**DOI:** 10.64898/2026.01.05.697699

**Authors:** Younes Valibeigi, Abouzar Kaboudian, Flavio Fenton, Gil Bub

**Affiliations:** Department of Physiology, McGill University, Montréal, QC, Canada; Montreal Neurological Institute, McGill University, Montréal, QC, Canada; School of Physics, Georgia Institute of Technology, Atlanta, GA, USA

**Author notes:** Corresponding author: Younes Valibeigi, Montreal Neurological Institute, McGill University, Montréal, QC, Canada.

**Keywords:** Digital Twin, cardiac electrophysiology, closed-loop systems, GPU computing, hybrid systems, optogenetics, reentrant arrhythmia

## Abstract

Reentrant arrhythmias are life-threatening cardiac events that are difficult to study due to limited experimental control over the complex circuit dynamics. We aimed to develop a real-time digital heart twin that allows in real-time dynamic manipulation of reentrant pathways in vitro using physiologically relevant simulations. We designed a closed-feedback loop system that couples a cultured cardiac monolayer with a two-dimensional computational simulation of cardiac tissue. The simulation, based on GPU-accelerated models (e.g., cellular automata), predicts wave propagation in real-time using the Abubu.js library. Optical mapping captures monolayer activation patterns, and simulation outputs are converted into light-based stimulation via optogenetics, using LEDs and microcontrollers to depolarize cardiac tissue. Our platform is capable of accurately detecting and responding to electrical waves in real-time, enabling interactive control of reentrant circuits. The system replaces traditional fixed-delay stimulation protocols with computationally guided interventions, better mimicking physiological conduction dynamics. This digital twin provides a novel and responsive method to study reentrant arrhythmias. Its integration of optical stimulation, real-time modeling, and tissue feedback enables the construction of user-defined reentry pathways under dynamic control. By merging computational and biological systems, this work introduces a versatile experimental framework for investigating arrhythmias. The platform may inform future control and anti-arrhythmic strategies and pave the way for personalized cardiac electrophysiology studies.

## I. Introduction

CARDIAC arrhythmias are disruptions in the normal initiation or propagation of electrical activity within the heart and are a major cause of mortality worldwide [1]. Among the various types of arrhythmias, those involving **reentrant circuits** are particularly important due to their role in life-threatening conditions such as ventricular tachycardia and fibrillation. Reentry occurs when an electrical wavefront continuously circulates within cardiac tissue, either around a fixed anatomical obstacle (*anatomical reentry*) or a functional pivot point, giving rise to spiral wave patterns (*functional reentry*) [2]. These spiral waves have been shown to exist in mammalian hearts [3], including human hearts [4], which can become unstable and multiply within the myocardium, eventually leading to fibrillation: a disorganized, unsynchronized state of excitation that impairs effective cardiac contraction and may result in sudden cardiac arrest [5,6].

Understanding the mechanisms by which reentrant waves form, stabilize, and transition into fibrillation is critical for advancing both diagnostic and therapeutic strategies. However, studying these dynamics experimentally is challenging due to the complexity and variability of cardiac tissue behavior. Closed-feedback loop systems, also known as digital twins, offer a powerful framework to address this challenge by coupling living cardiac tissue with computational simulations in real-time as it has been done with live neurons [7]. These systems allow researchers to impose dynamic and physiologically relevant perturbations based on feedback from the tissue itself, enabling precise control over reentrant circuits. By manipulating features such as circuit length, conduction velocity, and refractory period in a controlled manner, digital twins provide a unique opportunity to dissect the conditions that promote or suppress reentrant activity, and in particular, the transition to fibrillation.

A classic example of such a digital twin is the dynamic clamp, in which a computational model injects current into an excitable cell in response to its membrane voltage [8]. This approach has been successfully used in both neuronal [9] and cardiac [10] research. In cardiac electrophysiology, closed-loop digital twins have been developed to emulate and control reentrant circuits by modifying conduction parameters during cardiac activity [11]. Early implementations used analog and digital circuitry to achieve real-time responsiveness [12, 13], which evolved into software-based systems using real-time operating platforms such as RTXI [14] which was used to control in real-time discordant alternans in live Purkinje fibers [15].

A limitation of conventional stimulation methods in these systems is their reliance on electrodes, which are spatially static [15, 16] and do not require short distances to control even with advanced nonlinear control methods [17]. This constraint has led to the increasing use of optogenetics, which leverages light-sensitive ion channels expressed in genetically modified cardiac cells to achieve high-resolution, spatially flexible control of excitation patterns [18, 19]. Initial applications demonstrated the feasibility of using optogenetics for precise pacing and localization of pacemaker activity [20, 21]. Optical dynamic-clamp methods similarly showed real-time modulation of excitability with light-based feedback [22]. Subsequent studies applied spatial light patterns to modulate tissue behavior and terminate arrhythmias [23-25]. Further, all-optical systems combined optogenetic stimulation with simultaneous optical imaging of membrane voltage and intracellular calcium through a shared on-axis optical pathway [26]. Recent advances have demonstrated that micro-LED arrays can enable high-resolution control of cardiac monolayers through spatiotemporal optogenetic stimulation [27]. However, most of these implementations were open-loop, lacking real-time responsiveness to evolving tissue dynamics.

Recent approaches have explored fully optical closed-loop systems, where high-speed cameras are used to capture electrical activity and project stimulation patterns through digital light processors [11, 28]. However, these systems often depend on fixed-delay protocols, applying stimulation after a preset delay following tissue activation. Fixed-delay strategies assume linear conduction with constant velocity and ignore the complex, nonlinear properties of real cardiac tissue and complex spatial repolarization sequences. Such protocols cannot account for variability in wavefront shape, directional differences, or refractory behavior [11], and they typically stimulate the tissue at a fixed site, limiting spatial flexibility and failing to mimic dynamic wavefront interactions (Fig. 1(c)). More recently, opto-electronic feedback control of membrane potential has been introduced, enabling precise modulation of single-cell action potential waveforms in isolated cells or locally within multicellular preparations, although in these implementations the feedback is obtained from a single cell within the tissue [29].

**Fig. 1.**
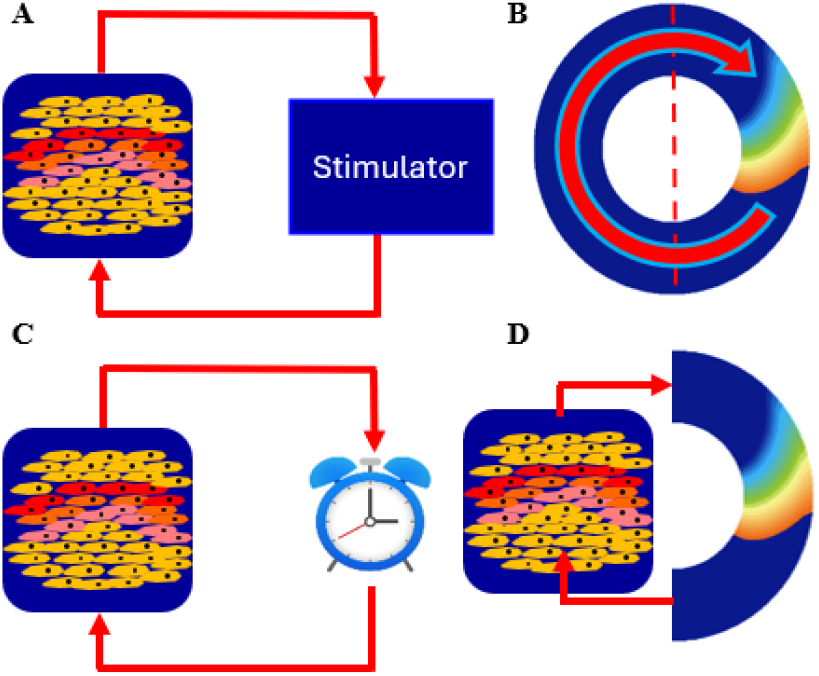
Conceptual overview of the closed-loop digital cardiac twin (A) Schematic of a closed-loop feedback system. (B) Cardiac simulation showing a reentrant wave circulating around an obstacle. (C) Closed-loop system incorporating a fixed delay. (D) Full implementation of the digital heart twin, where real-time signals from the tissue update the computational model, and model predictions guide adaptive control of reentrant activity.

Replacing fixed delays with real-time simulations that account for evolving tissue dynamics offers a more physiologically relevant strategy (Fig. 1(d)). However, traditional 2D cardiac models are computationally expensive and require dedicated workstations, making real-time interaction impractical with standard CPU-based simulations. To overcome this barrier, Kaboudian *et al*. [30] developed Abubu.js, a WebGL-based library that enables high-speed simulations of cardiac tissue on standard GPUs. This tool supports complex models such as Fenton-Karma [31] and OVVR [32], enabling dynamic wave propagation to be computed in real-time [33]. Crucially, it enables the generation of spatially and temporally variable reentrant wave patterns that are responsive to experimental feedback.

In this study, we introduce a fully optical, digital cardiac twin that integrates Abubu.js-based real-time cardiac simulations with optogenetically modified cardiac monolayers. By using matrix LEDs and microcontrollers to stimulate cardiac cells in response to both tissue activity and simulation output, our platform enables millisecond-scale bidirectional interaction. This setup allows for user-defined control over wavefront geometry and timing, offering a robust tool to explore how reentrant dynamics evolve and how they can be disrupted to control, prevent, and terminate fibrillation.

## II. Method

The digital twin establishes a bidirectional communication loop between the cardiac monolayer and a 2D cardiac simulation (or a fixed delay protocol), enabling real-time feedback control (Fig. 2). A Basler camera detects tissue excitations, which are processed in parallel by a simulation and relayed to an Arduino Uno microcontroller. The Arduino triggers patterned light stimulation via LEDs onto ChR2-expressing tissue. A NodeJS server coordinates data flow between components, maintaining synchronization between the camera and microcontroller for intervention within 2 ms of activity detection.

**Fig. 2.**
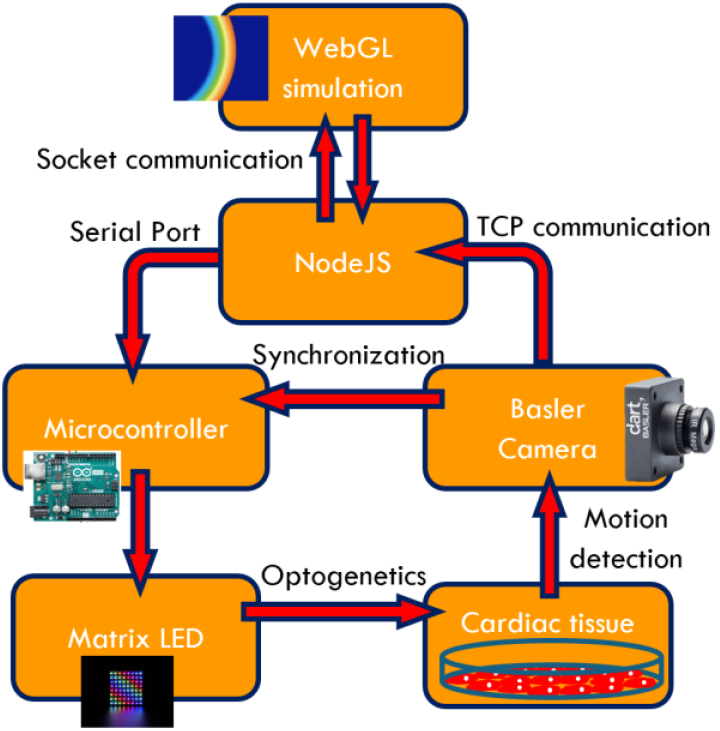
Overview of the closed-feedback loop system.

### A. Node.js Communication Architecture

Node.js is a back-end JavaScript runtime environment that enables server-side execution of JavaScript code outside the browser. In this system, Node.js functions as a communication hub, bridging the simulation interface with the camera, microcontroller (Arduino), and matrix LEDs through real-time sockets (Socket.IO), serial communication, and TCP protocols. A detailed description of the Node.js server setup and its communication architecture is available in the supplementary materials on the GitHub repository: https://github.com/younesvalibeigi/Digital-Heart-Twin.

### B. WebGL Simulation

To simulate excitation wave propagation in cardiac tissue, we used real-time 2D simulations implemented with Abubu.js, a JavaScript library built on WebGL for parallel GPU-based computation [30, 33, 34]. WebGL enables real-time modeling of large-scale cardiac systems by processing fragment and vertex shaders in parallel [35]. Abubu.js simplifies WebGL’s complex graphical pipeline, allowing computational variables to be stored and visualized using 4D textures. These simulations are embedded in an HTML environment and managed using JavaScript, where shaders receive and process data for each computational cell.

The server-side environment is managed using Node.js, which hosts the simulation and enables real-time communication with external hardware (e.g., microcontroller, camera) via Socket.IO and TCP protocols. The browser renders the WebGL simulation through a local server, and the main JavaScript file handles the transfer of data to shaders for visualization and control. A detailed breakdown of the Abubu.js simulation framework, folder structure, and Node.js communication architecture is available in the supplementary materials at: https://github.com/younesvalibeigi/Digital-Heart-Twin.

### C. Cardiac Simulations

We implemented a cardiac model using a cellular automaton (CA) algorithm [36, 37] within the Abubu.js library. The model simulates wave propagation in cardiac tissue, with real-time visualization enabled by GPU-accelerated WebGL rendering. Due to its simplicity, the CA approach supports rapid simulations while remaining capable of reproducing diverse physiological conditions, including unidirectional wave propagation in two-dimensional monolayers (Fig. 3(a)) and the generation of spiral wave dynamics (Fig. 3(b)) similar to the experimental monolayers. Implementation details, including shader logic and model parameters, are available in the supplementary materials (appendix A) and GitHub repository: https://github.com/younesvalibeigi/Digital-Heart-Twin.

**Fig. 3.**
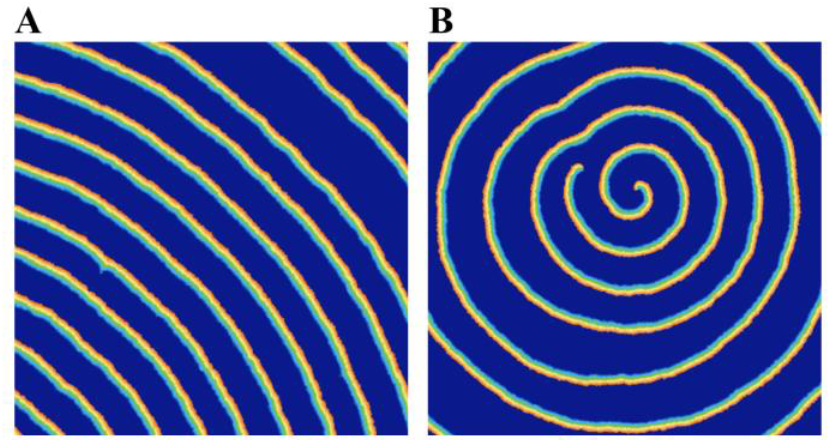
Two-dimensional cellular automata model implemented in WebGL using the Abubu.js library. (A) Simulation output showing the generation of uniform planar wavefronts propagating across the domain from the bottom left to the top right. (B) Simulation output illustrating the formation of spiral wave dynamics, demonstrating the model’s capability to reproduce complex reentrant patterns relevant to cardiac arrhythmias.

### D. Cardiac Monolayer Preparation and Viral Transduction

Postnatal mouse hearts (P0–P3) were isolated and enzymatically dissociated using a combination of extracellular matrix degradation and mechanical trituration to obtain a ventricular cell suspension. To minimize fibroblast contamination, cells were pre-plated on plastic for one hour before seeding. Monolayers of approximately 1 cm^2^ were cultured for recordings, with cell plating and media preparation protocols detailed in the supplementary materials, appendix B. Cultures were maintained at 37 °C with 5% CO_2_ in a humidified incubator, and recordings were conducted in a controlled environmental chamber under the same conditions. All procedures adhered to the Canadian Council on Animal Care guidelines and were approved under McGill protocol 2018-8044 (SOP 301-01).

Forty-eight hours after plating, ventricular monolayers were infected with adenovirus vector type 5 (dE1/D3) encoding channelrhodopsin-2 mutant H134R (ChR2(H134R)) at a multiplicity of infection (MOI) of 100. Culture media was refreshed every 48 hours post-infection. Additional details regarding infection and viral handling are provided in the supplementary materials, appendix B.

### E. Motion Detection Using High-Speed Imaging

A Basler Ace acA1920-155um camera operating at 30 frames per second (FPS) was used to capture optical signals from the cardiac monolayer. Camera configuration was performed using the Pylon 6.0.1 Camera Software Suite, with exposure time set to 30,000 µs and digital I/O settings configured for triggering on exposure active. The system recorded 500 frames per experiment, with each frame spanning 33.33 ms. Building on previous approaches [38, 39], we developed a motion-detection algorithm that identifies excitation waves from tissue displacement, rather than relying on traditional voltage-sensitive dye methods, thereby enabling cost-effective, long-duration recordings. Raw frames do not visibly reveal excitation waves; therefore, the algorithm computed the absolute difference between the current frame and its sixth predecessor. This operation highlights contractile activity as regions of intensity change, effectively visualizing propagating wavefronts across the tissue.

To achieve sub-frame temporal resolution with low-cost imaging, the algorithm analyzed motion across two regions of interest (ROIs) within the tissue. It first detected excitation in the first ROI by comparing pixel intensity averages and identifying the pixel column where the mean surpassed a predefined threshold. The same method was then applied to the second ROI. From the spatial and temporal difference between the two detections, the conduction velocity (CV) was calculated using the formula:

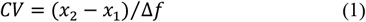

where *x*_1_ and *x*_2_ are the detected column positions and Δ*f* is the number of frames between detections. This allowed estimation of wave arrival time at specific regions based on velocity and distance:

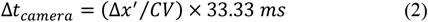

A full pseudocode and parameter settings for this algorithm are provided in the supplementary materials at: https://github.com/younesvalibeigi/Digital-Heart-Twin.

### F. Microcontroller and Matrix LED

Microcontrollers are compact, fast-processing units that integrate a CPU, clock, and I/O ports to digitally control external systems, including scientific instruments and simulate spiral waves [40]. In this work, an Arduino UNO was programmed using Arduino IDE (v1.8.49.0) to control an 8×8 Adafruit DotStar matrix LED. The microcontroller received buffer data via USB from the Node.js server through a serial port program. The matrix LED, soldered to the appropriate Arduino pins (GND, 5V, CLK to pin 13, and DIN to pin 11), projected blue light to activate ChR2-expressing cardiac cells. The simulation (main.js) can dynamically trigger customized LED patterns based on wave activity in real-time, enabling spatial and temporal optical control. A 3D base for the matrix LEDs was designed in SketchUp and printed using a Prusa MK2s printer. Implementation details and Arduino code are available in the supplementary materials on the GitHub page.

### G. Quantifying Communication Delays in the Closed-Loop Hardware Interface

To evaluate system accuracy, we used a photodiode (Thorlabs SM05PD1A) with a preamplifier (Thorlabs AMP 110), recorded by a digital oscilloscope (Picoscope) via the PicoLog 6 application. The photodiode captured the light output from the matrix LEDs with millisecond precision, registering each LED flash as a sharp drop in voltage. The intervals between successive flashes (cycle periods) were calculated and plotted in a histogram to assess timing consistency. A Python script for processing the photodiode data and generating histograms is available in the supplementary materials on the GitHub page. To identify sources of delay in the closed-loop system, we implemented three experimental designs. First, the Arduino was programmed to activate the LEDs periodically with a cycle of 1101 ms (this value was determined based on pilot experiments with the WebGL simulations) and demonstrated high temporal precision (mean cycle period = 1101.61 ms; standard deviation = 0.49 ms), confirming minimal delay in the microcontroller-LED interface. Second, periodic activation signals were sent to the Arduino via a Node.js script using serial port communication, which introduced increased variability (mean = 1105.66 ms; standard deviation = 5.62 ms), indicating latency within the communication pathway. Finally, the HTML client (running the WebGL simulation) was programmed to send periodic signals to the Node.js server, which then relayed them to the Arduino. This setup introduced additional delay and variability (mean = 1110.65 ms; standard deviation = 7.20 ms), attributed to the combined overhead of socket communication and browser execution. These findings demonstrated the need to synchronize the camera directly with the Arduino to bypass accumulated delays and preserve the temporal precision of real-time feedback.

### H. Synchronization of Camera and Microcontroller

To synchronize the Basler camera (acA1920-155um) with the Arduino microcontroller, a 6-wire Basler Power-I/O cable was used. The camera’s TTL output signal, generated at the start of each frame acquisition, was routed through the cable, with the brown wire connected to the Arduino’s analog input (A3) and the white wire to ground. This TTL pulse allowed the Arduino to count frames in real-time, effectively using the camera as a precise clock.

The camera software estimated the time required for a detected wave to reach a target region within the field of view (Δ*t*_*camera*_). This value was combined with the elapsed frame count (*F*) to calculate the total time since the beginning of acquisition: *t* = Δ*t*_*camera*_ + *F* × 33.33 *ms*. This timestamp was sent to the WebGL simulation, which modeled wave propagation across a virtual tissue domain. The simulation determined how many iterations (*F*_*simulation*_) were needed for a virtual wave to travel the simulated domain. This number was scaled to milliseconds using a conversion factor dt, derived from the domain length, estimated tissue conduction velocity (CV), and number of simulation iterations (Δt_simulation_ = *F*_*simulation*_ × *dt*).

The total time delay (*t*_*total*_ = Δ*t*_*camera*_ + Δ*t*_*simulation*_) was then sent via serial port communication to the Arduino. The Arduino converted this delay into a target camera frame and a residual delay (*t* = *F*_*total*_ × 33.33 *ms* + *Remainder*). It then waited for *F*_*total*_ TTL pulses from the camera and paused for the calculated remainder before activating the matrix LEDs. This approach eliminated the need for immediate microcontroller responses and bypassed variable delays introduced by socket.io or serial transmission, thereby improving the temporal accuracy of LED stimulation.

### I. Validation of System Timing Using Synthetic Wave Stimulation

To evaluate the system’s temporal precision, a DMD projector (Vialux XGA1303) was used to generate artificial excitation waves mimicking cardiac tissue dynamics. The projector was controlled by a computer running Jython software (a Python interpreter implemented in Java), which is available for download at: https://zenodo.org/records/5800192. Periodic wave patterns with 1100 ms intervals were projected onto the microscope stage. These optical signals were recorded by the Basler camera and processed by the WebGL simulation, which was configured to return a fixed delay rather than simulate wave propagation. After this programmed delay, the matrix LEDs were triggered using the feedback protocol described above. Since the projected waves had constant periodicity and the system applied a fixed delay, the cycle periods detected by the photodiode were expected to remain stable. This experimental setup provided a controlled means of assessing the accuracy and consistency of the device’s response timing.

### J. Optical Feedback Experimental System

The digital twin integrates real-time optical stimulation and imaging using a fully synchronized feedback loop. As shown in Fig. 6, the setup combines two microscope platforms (Olympus MVX10 and Nikon Eclipse Ti-U) to support wave imaging and optogenetic control, respectively [41]. A Basler acA1920-155um camera captures cardiac monolayer activity from above via the MVX10 microscope, while an 8×8 Adafruit DotStar LED matrix provides optogenetic stimulation from below through the Nikon Eclipse Ti-U’s optical path.

Node.js manages bidirectional communication between the WebGL-based simulation, the camera, and the Arduino Uno microcontroller. For validation experiments, a DMD projector projects spatially controlled artificial excitation waves onto the tissue plane. A photodiode coupled with a preamplifier records LED light output to verify timing precision. All components are linked via USB or TTL signal pathways, creating a closed-loop system capable of real-time detection and feedback actuation.

### III. Results

Here, we present a closed-feedback loop system that couples a cardiac monolayer with two-dimensional computational simulations, enabling dynamic modulation of wave propagation without relying on fixed-delay protocols. We demonstrate potential applications of this hybrid optical system for real-time control of cardiac monolayers.

### A. System Timing Accuracy and Sources of Temporal Error

Precise temporal synchronization is essential for digital cardiac twins, as even millisecond-scale deviations can alter the propagation dynamics of reentrant waves. In cultured cardiac monolayers, electrical wavefronts typically travel at CV = ∼10 μm/ms (comparable to those in iPSC-Cardiomyocyte monolayers[42]), making synchronization critical to avoid artifacts. We evaluated system accuracy by comparing the timing consistency of projected stimulation patterns with and without camera–microcontroller synchronization. The non-synchronized system exhibited a standard deviation of 11.34 ms. In contrast, synchronization reduced this deviation substantially, with a standard deviation of only 1.53 ms. This confirms that the synchronization protocol achieved the temporal resolution necessary for real-time cardiac feedback control.

Two principal factors influence timing precision: resolution error and signal noise. The camera’s spatial resolution (5.86 μm per pixel) limits the accuracy of wavefront position detection, with *x*_1_ ≈ 350 *pixels* and *x*_2_ ≈ 700 *pixels* (Fig. 4), each having an associated uncertainty of ±5.86 μm. This corresponds to a displacement of Δ*x* ≈ 350 *pixels*, or approximately 2051 μm. The combined spatial error in wave displacement, Δ*x*, is calculated as:

**Fig. 4.**
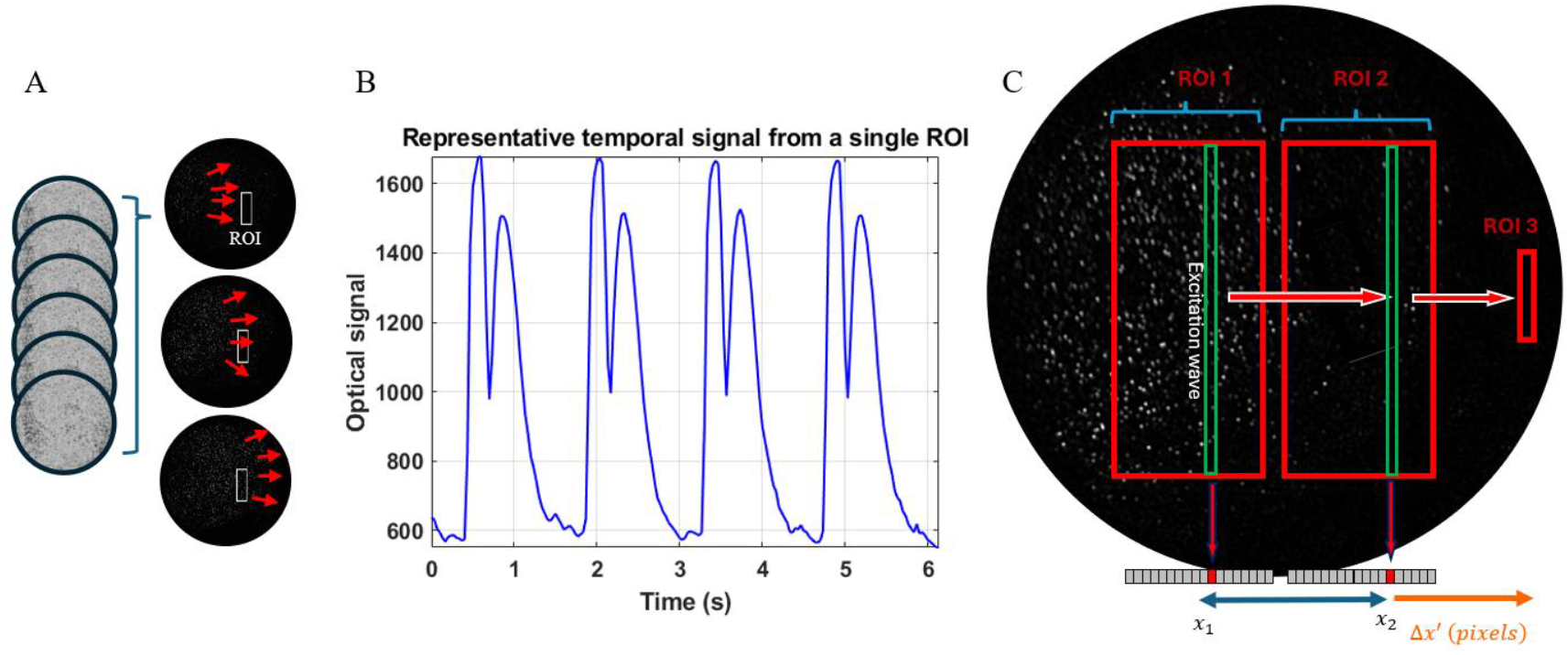
Motion-based wave detection algorithm. (A) Motion is extracted by subtracting the pixel intensity from its value six frames earlier, revealing tissue contractions and wave propagation. (B) Temporal signal from a representative region of interest (ROI, white rectangle) showing two intrinsic peaks corresponding to the contraction and relaxation phases of cardiac cells. (C) Excitation wave timing is determined by detecting peaks in pixel intensity across two regions of interest (ROI 1 and ROI 2). The algorithm calculates conduction velocity and estimates wave arrival time at ROI 3 based on spatial and temporal offsets. This value is transmitted to the feedback-loop controller.

**Fig. 5.**
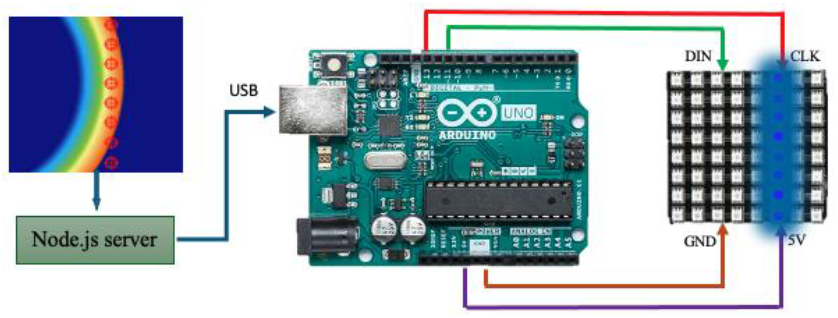
Real-time optogenetic stimulation via simulation-driven LED control. The simulation reads voltage values at specific coordinates in the 2D texture and, upon detecting activation, transmits a signal to the Node.js server. The server relays this information via serial communication to the Arduino microcontroller, which then triggers a defined pattern on the matrix LED. This enables synchronized light delivery to the cardiac monolayer based on simulated wave dynamics.

**Fig. 6.**
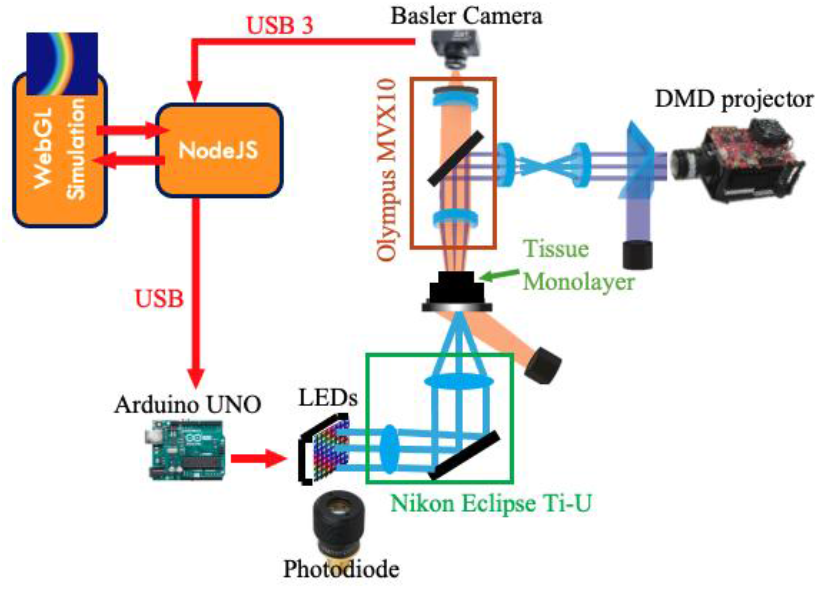
Schematic of the hybrid feedback system integrating imaging, optical stimulation, and simulation.

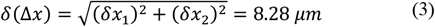

Given a displacement Δ*x*′ ≈ 2930 *μm* (Fig. 4) over 6 frames (*CV* = Δ*x*/Δ*F*), the conduction velocity (CV) uncertainty is:

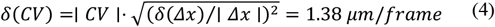

Consequently, the propagated error in timing, Δ*t*, based on frame rate (33.33 ms/frame), is:

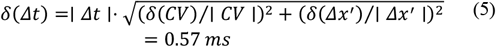

While this level of precision is sufficient for the experimental requirements, occasional deviations persisted even under synchronized conditions. As shown in Fig. 7, signal noise in the column-averaged pixel intensities used to detect wavefronts can introduce errors in peak detection. These signals inherently contain two peaks due to contraction and relaxation phases of cardiac cells. Small fluctuations or noise-induced deflections near threshold values can alter the detected timing by several milliseconds, leading to inaccurate estimations of wave velocity.

**Fig. 7.**
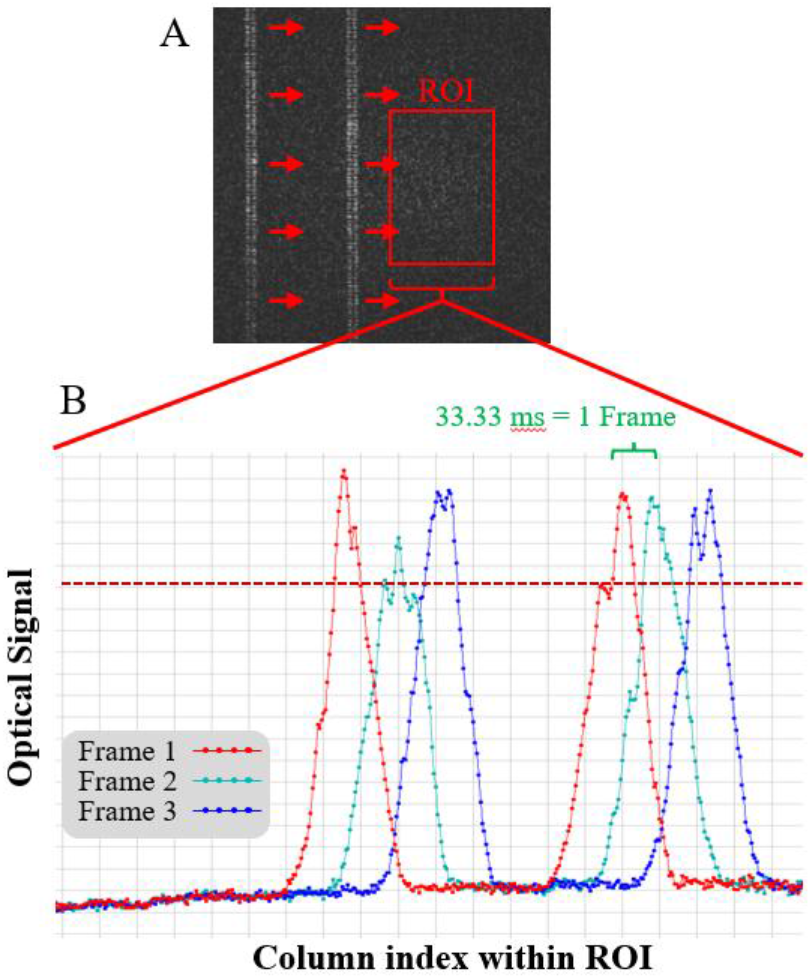
Source of inaccuracy due to signal noise. (A) A frame from simulated waves generated by the projector, demonstrating double peaks corresponding to the contraction and relaxation phases of virtual cardiac cells. (B) The plot shows the averaged optical signal across columns within the region of interest (ROI) for three consecutive frames. Due to the non-uniform shape of detected waves, increasing the detection threshold (dotted line) shifts the estimated timing of signal detection by several milliseconds, thereby introducing variability in wavefront localization and contributing to temporal noise within the system.

### B. Modulating Tissue Dynamics Using a Closed-Loop System with 2D Cellular Automata Simulation

Traditional closed-loop stimulation approaches for cardiac monolayers often rely on fixed-delay protocols [11] or simplified mathematical models [43]. In this study, we implemented a dynamic closed-feedback system that integrates a two-dimensional cellular automata simulation with a cardiac monolayer, enabling real-time bidirectional interaction. This hybrid configuration allowed us to manipulate tissue behavior based on computational model output, rather than fixed stimulation intervals.

As shown in Fig. 8, the model-driven stimulation modulated wave dynamics in the cardiac tissue. Specifically, comparison between the control condition (Fig. 8(a)) and the simulation-driven hybrid feedback condition (Fig. 8(b)) reveals alterations in wave propagation attributable to the system’s intervention. LED activation sites, indicating optogenetic stimulation, are visible, and column-averaged signals were derived from the defined regions of interest (white rectangles). Cycle length dynamics are plotted in Fig. 8(c), demonstrating alternating periods consistent with interaction between the simulation and the biological tissue, suggestive of emergent alternans behavior characterized by beat-to-beat alternation in electrical activity and contraction strength at a constant heart rate [44]. In a parallel experiment using a fixed-delay protocol (with 100 ms delays), shown in Fig. 8(e), the system similarly modulated tissue behavior, but with greater variability in wave dynamics, as quantified in Fig. 8(f). These results confirm that the digital twin can support real-time feedback using either delay-based or model-based control and reveal that model-based control can elicit distinct tissue dynamics compared to delay-based protocols, highlighting the system’s versatility for investigating complex reentrant phenomena.

**Fig. 8.**
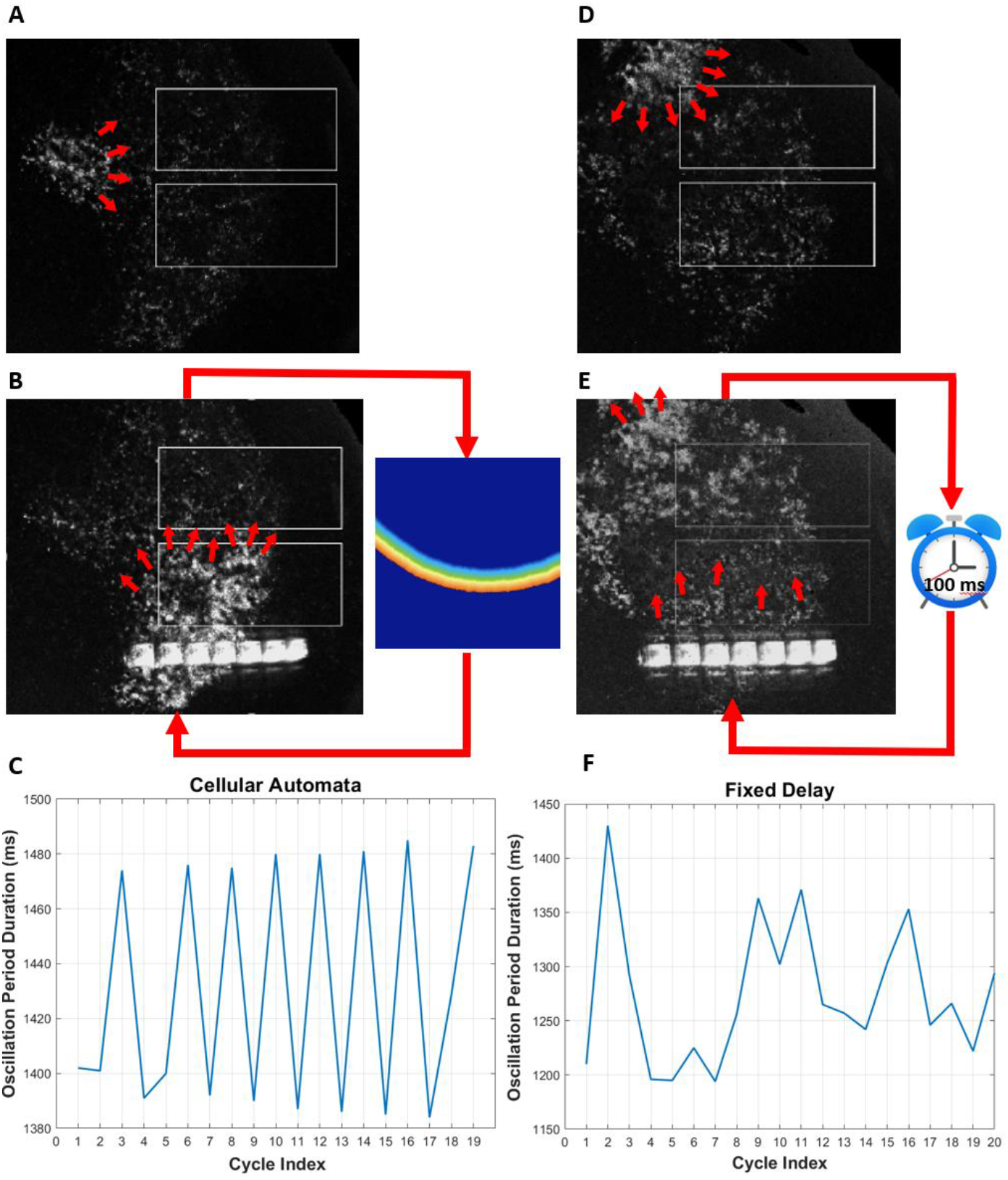
Digital twin with cellular automata model and fixed-delay protocol. (A) Cardiac activity recorded in the absence of model intervention (control). (B) Cardiac response under closed-loop stimulation with the cellular automata model. The two white rectangles indicate regions of interest positioned vertically to calculate wave propagation speed, assuming wavefronts travel from bottom to top. (C) Evolution of cycle periods during model-driven feedback, exhibiting an alternating pattern. (D) and (E) show a separate experimental sample under control and fixed-delay intervention, respectively. (F) Evolution of cycle periods under fixed-delay feedback, demonstrating irregular dynamics and lack of the periodicity observed with the cellular automata model.

## IV. Discussion

We developed a digital cardiac twin that links living monolayers with real-time 2D simulations using Abubu.js [33]. The system demonstrated closed-loop control, re-entrant wave generation, and potential for disease modeling, achieving millisecond precision with low-cost hardware. By replacing fixed delays with spatially extended simulations, this work introduces a novel and scalable approach for real-time cardiac modulation, with potential extension to 3D models, which can still be run in real-time using Abubu.js [45].

### A. Advantages over Previous Systems

Compared to earlier hybrid systems, the proposed setup introduces key improvements in accessibility, functionality, and integration. Prior approaches often relied on high-speed imaging equipment [11, 28, 43], which limits their adoption in resource-constrained environments. Our system addresses this by incorporating a custom algorithm capable of achieving high temporal accuracy using low-frame-rate machine vision cameras, thus significantly reducing experimental costs. In contrast to electrode-based stimulation techniques used in earlier studies [14, 46, 47], our fully optical, closed-loop system responds dynamically to spatially heterogeneous tissue activity and extends beyond single-cell feedback by integrating responses over large, multicellular regions of the tissue [29]. Unlike most cardiac optogenetic platforms that apply static stimuli regardless of real-time dynamics [24, 39], our design enables bidirectional interaction between the tissue and model. The system supports both fixed-delay protocols and dynamic 2D modeling, offering a versatile platform that overcomes previously outlined constraints of fixed-delay approaches [11, 28]. Finally, by relying on motion detection rather than voltage-sensitive dyes, our system is capable of long-duration experiments without phototoxic effects [48].

### B. Limitations of the digital heart twin

Despite these advancements, the digital twin has several limitations. First, the use of motion detection in place of voltage-based imaging may introduce inaccuracies. Although this dye-free approach enables longer experimental durations, it lacks the precision of voltage recordings and can misinterpret passive tissue motion as active wave propagation, particularly in low-contrast regions [48]. Second, the reliance on low-frame-rate cameras constrains temporal resolution. The system assumes wave concordance, i.e., constant propagation velocity across the tissue [49], which is not representative of discordant alternans, where velocity varies spatially. In such cases, the prediction algorithm fails to estimate accurate wave locations, limiting the scope of applicable experiments. Third, the system is sensitive to noise: tissue samples generating significant motion artifacts can degrade detection accuracy. Although future iterations may integrate Gaussian filtering to improve signal quality, this remains an unresolved challenge. Compared to high-speed camera-based systems that achieve precise real-time wave localization [11, 43], our algorithm estimates wave position using only two reference points. While this method works well for unidirectional and smoothly propagating waves, it is less effective when wavefronts change speed or direction. Moreover, our system has only been validated using cardiac monolayers, whereas other systems have demonstrated functionality in ex vivo intact hearts [11, 43]. Furthermore, the cellular automata model cannot generate electrophysiological waves with dynamic wavelengths and conduction velocity, in contrast to differential equation-based models such as the Fenton–Karma [31] and O’Hara–Virág– Varró–Rudy (OVVR) [32] models. This limitation restricts the system’s ability to reproduce certain physiological conditions associated with impaired ion conductance in cardiac cells, which the cellular automata framework cannot accurately simulate.

### C. Future Directions

This digital twin offers a powerful platform for studying the dynamics of re-entrant arrhythmias and other disruptive cardiac activities. Future developments include comparing fixed-delay protocols with 2D simulations to identify the most physiologically accurate modeling approach. Enhancing stimulation precision by replicating waveforms observed in simulations may improve the stability of re-entry and better mimic intact tissue behavior. Integrating AI-based computer vision for real-time detection and suppression of arrhythmic patterns could further expand the system’s therapeutic potential and support the development of adaptive, feedback-driven interventions.

Beyond the cellular automata framework used in this study, the platform can be implemented with more complex, differential equation-based cardiac models, such as the Fenton–Karma [31] or the OVVR models [32]. These models offer a richer set of physiological parameters for controlling cardiac dynamics and can generate wavefronts with variable wavelength and conduction velocity, enabling closer replication of intact tissue behavior [50]. Implementing such models within the GPU-accelerated Abubu.js framework could achieve simulation speeds comparable to real tissue activity, making real-time, bidirectional coupling with living cardiac monolayers feasible [33] and even possible with 3D hearts [45]. This capability would allow the system to reproduce a broad range of cardiovascular conditions and to support mechanistic studies of complex arrhythmias under precisely controlled experimental conditions.

## V. Conclusion

We developed a real-time digital cardiac twin that integrates living cardiac monolayers with GPU-accelerated 2D simulations to enable dynamic, closed-loop modulation of reentrant wave activity. This system introduces a novel framework that replaces fixed-delay protocols with spatially extended models capable of predicting wave propagation and interacting with biological tissue in real-time. Using a low-cost, accessible setup and custom algorithms, the system achieves millisecond-level precision without requiring high-speed imaging hardware, making it feasible for a broad range of research environments. It successfully demonstrated the generation and control of reentrant loops and holds potential for modeling pathological conduction conditions. By enabling bidirectional interaction between simulated and biological domains, the platform offers a versatile tool for studying arrhythmic behaviors, identifying the mechanisms that sustain them, and evaluating dynamic intervention strategies. The dye-free optical configuration supports long-duration experiments, providing opportunities for chronic studies and therapeutic development. Looking ahead, the digital twin can be expanded with more detailed simulations, 3D modeling, and real-time AI-based detection to enhance its clinical applicability. Ultimately, this work lays the foundation for adaptive, patient-specific interventions by coupling computational modeling with live cardiac tissue in a responsive and scalable framework.

## Appendix

A detailed explanation of the cellular automaton model is provided in Appendix A of the supplementary materials. The procedures for monolayer tissue preparation are described in Appendix B. The full code implementation for each component of the digital twin, along with comprehensive documentation and explanations, is available on GitHub: https://github.com/younesvalibeigi/Digital-Heart-Twin.

## References

[1] D. P. Zipes, and J. Jalife, Cardiac electrophysiology : from cell to bedside, 8th ed., Philadelphia, PA: Elsevier, 2021.

[2] E. M. Cherry, and F. H. Fenton, “Visualization of spiral and scroll waves in simulated and experimental cardiac tissue,” New journal of physics, vol. 10, no. 12, pp. 125016, 2008.

[3] A. M. Pertsov, J. M. Davidenko, R. Salomonsz, W. T. Baxter, and J. Jalife, “Spiral waves of excitation underlie reentrant activity in isolated cardiac muscle,” Circulation Research, vol. 72, no. 3, pp. 631–650, 1993.

[4] I. Uzelac, S. Iravanian, N. K. Bhatia, and F. H. Fenton, “Direct observation of a stable spiral wave reentry in ventricles of a whole human heart using optical mapping for voltage and calcium,” Heart Rhythm, vol. 19, no. 11, pp. 1912–1913, Nov, 2022.

[5] A. V. Panfilov, “Spiral breakup as a model of ventricular fibrillation,” Chaos: An Interdisciplinary Journal of Nonlinear Science, vol. 8, no. 1, pp. 57–64, 1998/03/01/, 1998.

[6] I. Uzelac, S. Iravanian, N. K. Bhatia, and F. H. Fenton, “Spiral wave breakup: Optical mapping in an explanted human heart shows the transition from ventricular tachycardia to ventricular fibrillation and self-termination,” Heart Rhythm, vol. 19, no. 11, pp. 1914–1915, Nov, 2022.

[7] T. J. Kispersky, M. N. Economo, P. Randeria, and J. A. White, “GenNet: A Platform for Hybrid Network Experiments,” Frontiers in Neuroinformatics, vol. 0, 2011, 2011.

[8] A. A. Prinz, and R. H. Cudmore, “Dynamic clamp,” Scholarpedia, vol. 6, no. 5, pp. 1470, 2011/05/02/, 2011.

[9] A. A. Sharp, M. B. O’Neil, L. F. Abbott, and E. Marder, “Dynamic clamp: computer-generated conductances in real neurons,” Journal of Neurophysiology, vol. 69, no. 3, pp. 992–995, 1993/03/01/, 1993.

[10] R. Wilders, “Dynamic clamp: a powerful tool in cardiac electrophysiology,” The Journal of Physiology, vol. 576, no. 2, pp. 349–359, 2006, 2006.

[11] M. Scardigli, C. Müllenbroich, E. Margoni, S. Cannazzaro, C. Crocini, C. Ferrantini, R. Coppini, P. Yan, L. M. Loew, M. Campione, L. Bocchi, D. Giulietti, E. Cerbai, C. Poggesi, G. Bub, F. S. Pavone, and L. Sacconi, “Real-time optical manipulation of cardiac conduction in intact hearts,” J Physiol, vol. 596, no. 17, pp. 3841–3858, Sep, 2018.

[12] A. A. Sharp, L. F. Abbott, and E. Marder, “Artificial electrical synapses in oscillatory networks,” Journal of Neurophysiology, vol. 67, no. 6, pp. 1691–1694, 1992/06/01/, 1992.

[13] H. P. C. Robinson, and N. Kawai, “Injection of digitally synthesized synaptic conductance transients to measure the integrative properties of neurons,” Journal of Neuroscience Methods, vol. 49, no. 3, pp. 157–165, 1993/09/01/, 1993.

[14] Y. A. Patel, A. George, A. D. Dorval, J. A. White, D. J. Christini, and R. J. Butera, “Hard real-time closed-loop electrophysiology with the Real-Time eXperiment Interface (RTXI),” PLoS Comput Biol, vol. 13, no. 5, pp. e1005430, May, 2017.

[15] D. J. Christini, M. L. Riccio, C. A. Culianu, J. J. Fox, A. Karma, and R. F. Gilmour, “Control of Electrical Alternans in Canine Cardiac Purkinje Fibers,” Physical Review Letters, vol. 96, no. 10, pp. 104101, 03/17/, 2006.

[16] W.-J. Rappel, F. Fenton, and A. Karma, “Spatiotemporal Control of Wave Instabilities in Cardiac Tissue,” Physical Review Letters, vol. 83, no. 2, pp. 456–459, 07/12/, 1999.

[17] A. Garzón, R. O. Grigoriev, and F. H. Fenton, “Model-based control of cardiac alternans in Purkinje fibers,” Physical Review E, vol. 84, no. 4, pp. 041927, 10/21/, 2011.

[18] E. Entcheva, and G. Bub, “All-optical control of cardiac excitation: combined high-resolution optogenetic actuation and optical mapping,” The Journal of Physiology, vol. 594, no. 9, pp. 2503–2510, 2016, 2016.

[19] V. Emiliani, E. Entcheva, R. Hedrich, P. Hegemann, K. R. Konrad, C. Lüscher, M. Mahn, Z.-H. Pan, R. R. Sims, J. Vierock, and O. Yizhar, “Optogenetics for light control of biological systems,” Nature Reviews Methods Primers, vol. 2, no. 1, pp. 55, 2022/07/21, 2022.

[20] A. B. Arrenberg, D. Y. R. Stainier, H. Baier, and J. Huisken, “Optogenetic control of cardiac function,” Science (New York, N.Y.), vol. 330, no. 6006, pp. 971–974, 2010/11/12/, 2010.

[21] T. Bruegmann, D. Malan, M. Hesse, T. Beiert, C. J. Fuegemann, B. K. Fleischmann, and P. Sasse, “Optogenetic control of heart muscle in vitro and in vivo,” Nature Methods, vol. 7, no. 11, pp. 897–900, 2010/11//, 2010.

[22] B. Quach, T. Krogh-Madsen, E. Entcheva, and D. J. Christini, “Light-Activated Dynamic Clamp Using iPSC-Derived Cardiomyocytes,” Biophysical Journal, vol. 115, no. 11, pp. 2206–2217, 2018.

[23] T. Bruegmann, P. M. Boyle, C. C. Vogt, T. V. Karathanos, H. J. Arevalo, B. K. Fleischmann, N. A. Trayanova, and P. Sasse, “Optogenetic defibrillation terminates ventricular arrhythmia in mouse hearts and human simulations,” The Journal of Clinical Investigation, vol. 126, no. 10, pp. 3894–3904, 2016/10/03/, 2016.

[24] C. Crocini, C. Ferrantini, R. Coppini, M. Scardigli, P. Yan, L. M. Loew, G. Smith, E. Cerbai, C. Poggesi, F. S. Pavone, and L. Sacconi, “Optogenetics design of mechanistically-based stimulation patterns for cardiac defibrillation,” Scientific Reports, vol. 6, no. 1, pp. 35628, 2016/10/17/, 2016.

[25] E. C. A. Nyns, A. Kip, C. I. Bart, J. J. Plomp, K. Zeppenfeld, M. J. Schalij, A. A. F. de Vries, and D. A. Pijnappels, “Optogenetic termination of ventricular arrhythmias in the whole heart: towards biological cardiac rhythm management,” European Heart Journal, vol. 38, no. 27, pp. 2132–2136, 2017/07/14/, 2017.

[26] A. Klimas, G. Ortiz, S. C. Boggess, E. W. Miller, and E. Entcheva, “Multimodal on-axis platform for all-optical electrophysiology with near-infrared probes in human stem-cell-derived cardiomyocytes,” Progress in Biophysics and Molecular Biology, vol. 154, pp. 62–70, 2020/08/01/, 2020.

[27] S. Junge, M. E. Ricci Signorini, M. Al Masri, J. Gülink, H. Brüning, L. Kasperek, M. Szepes, M. Bakar, I. Gruh, A. Heisterkamp, and M. L. Torres-Mapa, “A micro-LED array based platform for spatio-temporal optogenetic control of various cardiac models,” Scientific Reports, vol. 13, no. 1, pp. 19490, 2023/11/09, 2023.

[28] V. Biasci, L. Sacconi, E. N. Cytrynbaum, D. A. Pijnappels, T. De Coster, A. Shrier, L. Glass, and G. Bub, “Universal mechanisms for self-termination of rapid cardiac rhythm,” Chaos: An Interdisciplinary Journal of Nonlinear Science, vol. 30, no. 12, pp. 121107, 2020/12/01/, 2020.

[29] B. Ördög, T. De Coster, S. O. Dekker, C. I. Bart, J. Zhang, G. J. J. Boink, W. H. Bax, S. Deng, B. L. den Ouden, A. A. F. de Vries, and D. A. Pijnappels, “Opto-electronic feedback control of membrane potential for real-time control of action potentials,” Cell Reports Methods, vol. 3, no. 12, 2023.

[30] A. Kaboudian, E. M. Cherry, and F. H. Fenton, “Real-time interactive simulations of large-scale systems on personal computers and cell phones: Toward patient-specific heart modeling and other applications,” Science Advances, vol. 5, no. 3, pp. eaav6019, 2019.

[31] F. Fenton, and A. Karma, “Vortex dynamics in three-dimensional continuous myocardium with fiber rotation: Filament instability and fibrillation,” Chaos: An Interdisciplinary Journal of Nonlinear Science, vol. 8, no. 1, pp. 20–47, 1998/03/01/, 1998.

[32] T. O’Hara, L. Virág, A. Varró, and Y. Rudy, “Simulation of the Undiseased Human Cardiac Ventricular Action Potential: Model Formulation and Experimental Validation,” PLOS Computational Biology, vol. 7, no. 5, pp. e1002061, 2011.

[33] A. Kaboudian, E. M. Cherry, and F. H. Fenton, “Large-scale Interactive Numerical Experiments of Chaos, Solitons and Fractals in Real Time via GPU in a Web Browser,” Chaos Solitons Fractals, vol. 121, pp. 6–29, Apr, 2019.

[34] A. Kaboudian, “kaboudian/abubujs,” 2021.

[35] F. L. Patricio Gonzalez Vivo. “The Book of Shaders,” https://thebookofshaders.com/.

[36] H. Zhu, P. Y. Pang, Y. Sun, and P. Dhar, “Asynchronous adaptive time step in quantitative cellular automata modeling,” BMC Bioinformatics, vol. 5, pp. 85, Jun 29, 2004.

[37] G. Bub, A. Shrier, and L. Glass, “Spiral wave generation in heterogeneous excitable media,” Physical Review Letters, vol. 88, no. 5, pp. 058101, 2002/02/04/, 2002.

[38] L. Sala, B. J. van Meer, L. G. J. Tertoolen, J. Bakkers, M. Bellin, R. P. Davis, C. Denning, M. A. E. Dieben, T. Eschenhagen, E. Giacomelli, C. Grandela, A. Hansen, E. R. Holman, M. R. M. Jongbloed, S. M. Kamel, C. D. Koopman, Q. Lachaud, I. Mannhardt, M. P. H. Mol, D. Mosqueira, V. V. Orlova, R. Passier, M. C. Ribeiro, U. Saleem, G. L. Smith, F. L. Burton, and C. L. Mummery, “MUSCLEMOTION,” Circulation Research, vol. 122, no. 3, pp. e5–e16, 2018.

[39] G. Bub, and R.-A. B. Burton, “Macro-micro imaging of cardiac– neural circuits in co-cultures from normal and diseased hearts,” The Journal of Physiology, vol. 593, no. 14, pp. 3047–3053, 2015, 2015.

[40] A. J. Welsh, C. Delgado, C. Lee-Trimble, A. Kaboudian, and F. H. Fenton, “Simulating waves, chaos and synchronization with a microcontroller,” Chaos: An Interdisciplinary Journal of Nonlinear Science, vol. 29, no. 12, pp. 123104, 2019/12/01/, 2019.

[41] R. A. B. Burton, A. Klimas, C. M. Ambrosi, J. Tomek, A. Corbett, E. Entcheva, and G. Bub, “Optical control of excitation waves in cardiac tissue,” Nature Photonics, vol. 9, no. 12, pp. 813–816, 2015/12//, 2015.

[42] W. Dou, Q. Zhao, M. Malhi, X. Liu, Z. Zhang, L. Wang, S. Masse, K. Nanthakumar, R. Hamilton, J. T. Maynes, and Y. Sun, “Label-free conduction velocity mapping and gap junction assessment of functional iPSC-Cardiomyocyte monolayers,” Biosensors and Bioelectronics, vol. 167, pp. 112468, 2020/11/01/, 2020.

[43] S. Iravanian, and D. J. Christini, “Optical mapping system with real-time control capability,” American Journal of Physiology-Heart and Circulatory Physiology, vol. 293, no. 4, pp. H2605–H2611, 2007/10/01/, 2007.

[44] J. N. Edwards, and L. A. Blatter, “Cardiac alternans and intracellular calcium cycling,” Clin Exp Pharmacol Physiol, vol. 41, no. 7, pp. 524–32, Jul, 2014.

[45] A. Kaboudian, R. A. Gray, I. Uzelac, E. M. Cherry, and F. H. Fenton, “Fast interactive simulations of cardiac electrical activity in anatomically accurate heart structures by compressing sparse uniform cartesian grids,” Computer Methods and Programs in Biomedicine, vol. 257, pp. 108456, 2024/12/01/, 2024.

[46] D. J. Christini, and J. J. Collins, “Using chaos control and tracking to suppress a pathological nonchaotic rhythm in a cardiac model,” Physical Review E, vol. 53, no. 1, pp. R49–R52, 1996/01/01/, 1996.

[47] L. H. Frame, and M. B. Simson, “Oscillations of conduction, action potential duration, and refractoriness. A mechanism for spontaneous termination of reentrant tachycardias,” Circulation, vol. 78, no. 5 Pt 1, pp. 1277–1287, 1988/11//, 1988.

[48] J. Christoph, M. Chebbok, C. Richter, J. Schröder-Schetelig, P. Bittihn, S. Stein, I. Uzelac, F. H. Fenton, G. Hasenfuß, R. F. Gilmour Jr, and S. Luther, “Electromechanical vortex filaments during cardiac fibrillation,” Nature, vol. 555, no. 7698, pp. 667–672, 2018/03//, 2018.

[49] R. H. Anderson, E. A. Shinebourne, and L. M. Gerlis, “Criss-Cross Atrioventricular Relationships Producing Paradoxical Atrioventricular Concordance or Discordance,” Circulation, vol. 50, no. 1, pp. 176–180, 1974/07/01/, 1974.

[50] E. G. Tolkacheva, D. G. Schaeffer, D. J. Gauthier, and C. C. Mitchell, “Analysis of the Fenton–Karma model through an approximation by a one-dimensional map,” Chaos: An Interdisciplinary Journal of Nonlinear Science, vol. 12, no. 4, pp. 1034–1042, 2002/12/01/, 2002.

