## Supplementary Materials for "Real-Time GPU-Accelerated Digital Heart Twin: Integrating Bidirectional Interactions Between Living Optogenetic Monolayers and Computational Simulations"

### Appendix A. Implementation of Cardiac Models in AbubuJS

#### Overview

This document provides detailed implementation information for the cardiac simulation model used in the hybrid system: the Cellular Automata (CA) model. The model is implemented using the AbubuJS library for GPU-accelerated real-time simulation via WebGL shaders.

#### A1. Cellular Automata (CA) Model

##### A1.1 Theoretical Background

Cellular automata (CA) are discrete computational models that simulate the propagation of excitation waves using simplified rules based on cell states and neighborhood interactions. CA models are computationally efficient and particularly suited for modeling large-scale tissue dynamics with minimal computational cost.

In cardiac CA, each cell exists in one of three basic states:

- **Excitable** (voltage < 0.05)
- **Active** (voltage = 0.98)
- **Refractory** (0.05 ≤ voltage < 0.98)

Cells are activated based on a threshold ratio of active neighboring cells within a defined radius. After activation, a refractory period prevents immediate reactivation.

##### A1.2 Model Parameters

- **Excitability radius**: Number of grid units to search around each cell (e.g., 5–10 units)
- **Threshold**: Ratio of active neighbors required for activation (e.g., 0.3–0.4)
- **Refractory decrement**: Voltage decreases by 0.051 per time step

##### A1.3 Simulation Initialization

All cells are initialized with zero voltage. An external trigger or user click initiates excitation in a region of the grid.

##### A1.4 Shader Logic

At each iteration, every cell evaluates the number of active neighboring cells (defined as those with voltage < 0.98) within a specified radius (Fig. 1A). If the proportion of active cells within this radius exceeds a predefined threshold, the cell transitions to the active state (voltage = 0.98) in the subsequent iteration. Cells with voltage values greater than 0.05 are considered to be in the recovery phase; these cells are refractory and cannot be reactivated. Their voltage decreases incrementally at each time step by a fixed value (ΔV = 0.051) until they return to the resting state. The fragment shader (GLSL) implements the following logic for each cell:

```
if (voltage < 0.05) { // Cell is excitable
```

```
    if (active_neighbor_ratio > threshold) {
        voltage = 0.98; // Activate cell
    }
} else {
    voltage -= 0.051; // Recovery phase
}
```

Each simulation step involves two render passes to alternate between textures and allow iterative updating. The full implementation is available at: https://github.com/younesvalibeigi/Hybrid-Cardiac-Model. Fig. 1 illustrates how the implemented algorithm enables wave propagation across the tissue.

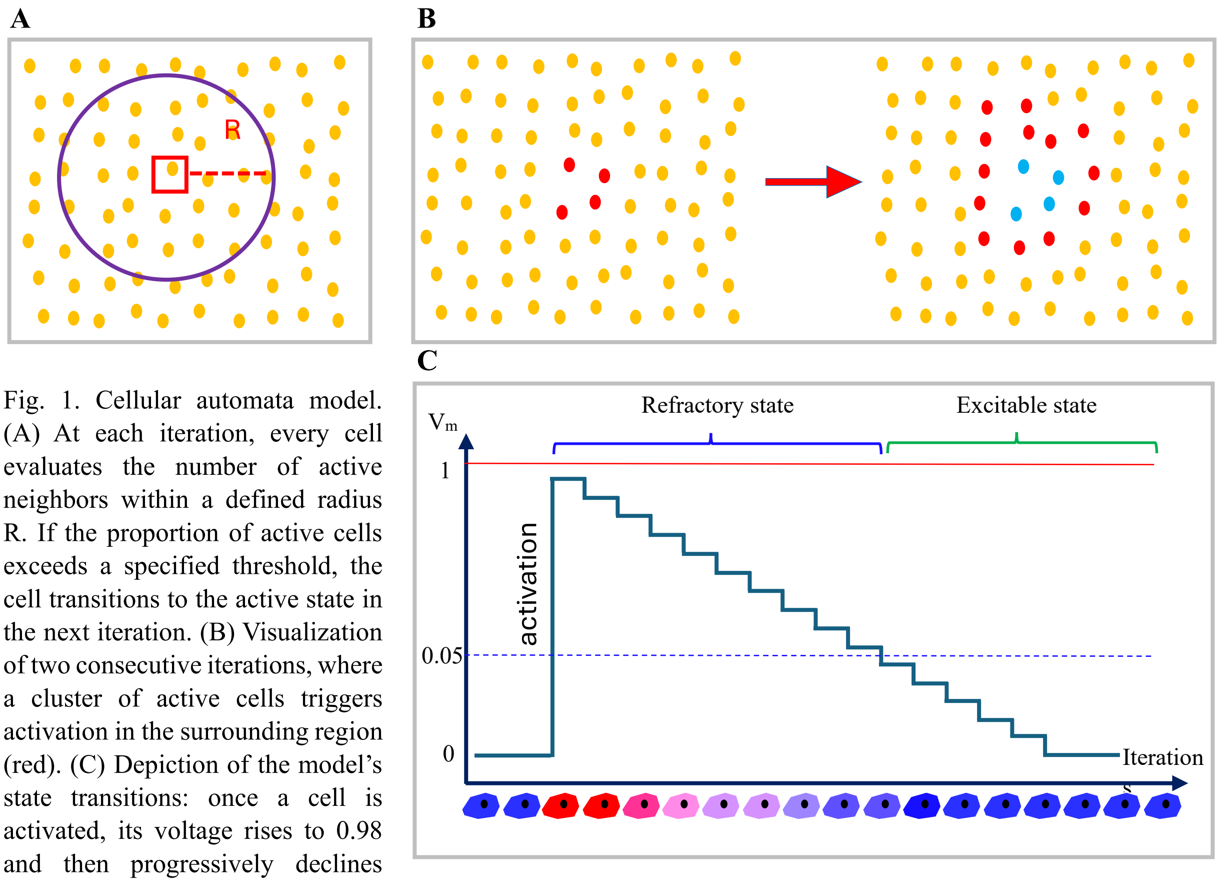


Fig. 1. Cellular automata model. (A) At each iteration, every cell evaluates the number of active neighbors within a defined radius R. If the proportion of active cells exceeds a specified threshold, the cell transitions to the active state in the next iteration. (B) Visualization of two consecutive iterations, where a cluster of active cells triggers activation in the surrounding region (red). (C) Depiction of the model's state transitions: once a cell is activated, its voltage rises to 0.98 and then progressively declines with each subsequent iteration, representing the refractory phase.

#### A1.5 Performance

The CA model runs at real-time speeds on standard GPUs (e.g., 60 FPS on NVIDIA GTX 1050) and is suitable for high-resolution grids (e.g., 256×256).

#### A2. Access to Code

The complete source code for both models, including GLSL shaders, main.js, and configuration files, is available at:

https://github.com/younesvalibeigi/Hybrid-Cardiac-Model

Implementation examples and documentation are also provided to assist new users in customizing the simulation for their own research.

### Appendix B. Monolayer preparation

The following protocol follows a step-by-step procedure from Miltenyi Biotec (Protocol No. 130-098-373, Miltenyi Biotec Inc, California, US), with additional steps from (Ambrosi et al., 2014; Burton et al., 2015). The entire protocol was first described in a submitted MSc thesis from a student in the same laboratory (Sepúlveda, 2020).

Cells were isolated using a Miltenyi gentleMACS Dissociator, which automates the cell dissociation process. Procedures for animal handling were performed in agreement with guidelines of the Canadian Council on Animal Care. Neonates were euthanized by decapitation in agreement with McGill University SOP 301-01 under approved protocol 2018-8044. Sterile techniques are followed during all procedures.

#### B.1. Tissue culture coating (for 24-well plates) and plating cell densities:

1. 5 µL of fibronectin (VWR, Ontario, CA) is diluted into 500 µL of PBS for a final concentration of 50 µg/µL.
2. 125 µL of thoroughly mixed dilution must be added to each well.
3. The plate should be incubated at 37 ºC and 5% CO2 for at least 1 h before use.

The densities described in the table below were used for neonatal mice cardiomyocytes.

| Cell density and seeding volume (Table F1) | | | |
|---|---|---|---|
| Vessel size | Surface area in $cm^2$ | Seeding volume in ml | Cell number (~156 x $10^3$ cells/$cm^2$) |
| Glass ring | 1 | 0.232 | 156 x $10^3$ |
| 24-well | 1.9 | 0.6 | 296 x $10^3$ |
| 12-well | 3.8 | 1.2 | 593 x $10^3$ |

#### B.2. Ventricle dissociation, tissue culture, and adenoviral infection of cardiac monolayers:

i. Prepare necessary solutions from stocks Penicillin/Streptomycin Mixture (Pen/Strep, Quality Biological, Cat#: 120-095-721), Dulbecco's Modified Eagle Medium (DMEM,

Wisent, REF: 319-062-CL), Phosphate Buffered Saline (PBS, Wisent, REF: 311-010-CL), and Fetal Bovine Serum (FBS, Gibco, Cat#12483020). DMEM (10%) maintenance media is prepared by adding 50 mL of FBS and 5 mL on Pen/Strep to 450 mL of DMEM, and DMEM (2%) is prepared by adding 200 µL of FBS and 100 µL of Pen/Strep to 10mL of DMEM.

ii. Harvest the heart and dissect the ventricles:
   a. Obtain a litter (6 or more animals) of postnatal 0 – 3 (P0 - P3) day old mice pups.
   b. Pups are decapitated, and the heart is isolated by first opening the rib cage with sharp scissors and remove the heart with tweezers. The hearts are placed in a shallow plate containing 3 ml of PBS solution on ice.
   c. Scissors are used to remove the ventricles (around the lower 70% portion of the heart) and remaining connective tissue.
   d. The tissue is cleaned by swirling the plate regularly to remove blood cells from the ventricles.
   e. The ventricles are each cut into 4 – 6 small pieces around 1–2 $mm^3$.

iii. The ventricles are dissociated into single cells using the Neonatal Heart Dissociation Kit and gentleMACS Dissociator, following protocol 130-098-373. The program used on the dissociator was "m_neoheart_01_01". The protocol takes approximately 45 minutes.
   a. The resulting suspension is centrifuged at 600xg for 5 mins.
   b. The pellet is resuspended in 10 mL of DMEM 10% using gentle manual agitation with a wide-mouthed pipette.

iv. Remove cardiac fibroblasts and count cells:
   a. The 10 mL suspension is transferred to a shallow 10 cm culture dish and incubated (at 37 degrees, 5% CO2) for 45 minutes. Fibroblasts settle and adhere to the bottom of the dish. The supernatant contains an enriched population of myocytes.
   b. The supernatant (10ml) and an additional 5ml PBS for washing the plate are transferred to a 50ml Falcon tube.
   c. Centrifuge the 15ml supernatant at 600xg for 5 min and resuspend the pellet in 1 mL of warmed DMEM 10%.
   d. Cell concentration is determined by using a standard hemocytometer, with Trypan Blue added to count dead cells.

v. Culture cardiomyocyte cells in 24-well plates:
   a. Once counted, the desired concentration per well is determined by table F1 above (approximately 300,000 cells per well in a 24 well plate).
   b. An additional 1 mL of DMEM 10% per well is added, and the tissue incubated for 24 hours.
   c. Provide post-plating maintenance by replacing the solution with 2 mL of DMEM 10% every 24-48 hours.

vi. Infect the ventricular myocytes by Ad-CMV-hChR2(H134R)-eYFP (Ambrosi et al., 2014) 2 to 3 days post plating:
   a. Replace DMEM 10% with a small amount of DMEM 2% (250 µL per plate).
   b. For our desired multiplicity of infection (MOI) of 100 and a virus titer of 2.2x106, we use 13.6 µL of virus solution per well (assuming 300,000 cells/well).
   c. Incubate monolayers for 48 h prior to running experiments

#### B3. External References


1. Ambrosi, C.M. and E. Entcheva, *Optogenetic Control of Cardiomyocytes via Viral Delivery.* Methods in molecular biology (Clifton, N.J.), 2014. **1181**: p. 215-228.
2. Burton, R.A.B., et al., *Optical control of excitation waves in cardiac tissue.* Nature Photonics, 2015. **9**(12): p. 813-816.
3. Sepúlveda, J.R., *Optically induced heterogeneities in cardiac tissue*. 2020, McGill University: Montreal. p. 76.
